# Proteome dynamics of COVID-19 severity learnt by a graph convolutional network of multi-scale topology

**DOI:** 10.1101/2022.07.04.498661

**Authors:** Samy Gauthier, Alexy Tran-Dinh, Ian Morilla

## Abstract

Many efforts have been recently done to characterise the molecular mechanisms of COVID-19 disease. These efforts resulted in a full structural identification of ACE2 as principal receptor of the Sars-CoV-2 spike protein in the cell. However, there are still important open questions related to other proteins involved in the progression of the disease. To this end, we have modelled the plasma proteome of 384 COVID patients. The model calibrated proteins measures at three time tags and make also use of the detailed clinical evaluation outcome of each patient after their hospital stay at day 28. Our analysis is able to discriminate severity of the disease by means of a metric based on available WHO scores of disease progression. Then, we identify by topological vectorisation those proteins shifting the most in their expression depending on that severity classification. Finally, the extracted topological invariants respect the protein expression at different times were used as base of a graph convolutional network. This model enabled the dynamical learning of the molecular interactions produced between the identified proteins.

## Introduction

There exist proved evidences on how certain proteins such as ACE2 receptor or TMPRSS2 are used by severe acute respiratory syndrome coronavirus 2 (Sars-Cov-2) as entrance gates to infect the cell. Likewise, there are also multiple clues on a likely participation of other proteins in the downstream of the disease during its progression Delgado Blanco et al. 2020; Scudellari 2021; Yang, Petitjean, and Koehler 2020; Zamorano Cuervo and Grandvaux 2020. All these experimental efforts aim to characterise the progression of COVID from an in situ baseline analysis of proteomic profiles. Unfortunately, most of those analyses tend to overlooking non-linear programmes of interaction determined by subsets of proteins already described in such studies. Some of those programmes are merely contributing to an innocuous reconfiguration of the secondary immune response system, but others can be causally provoking a worsening in the severity of the symptoms during the disease progression. In this work, we learn the later through graph convolutional networks calibrated with higher topological features extracted from raw soluble proteomic data. To this end, we computed persistent homology of more than 1, 400 protein profiles in blood enhanced in endothelial cells across samples Xu et al. 2019. Our models achieved tracking protein interactions occurring post-infection by which different levels of severity developed by 384 individuals suffering from Covid19 symptoms might be explained Filbin, Goldberg, and Hacohen cited October 2021. In this sense, the Covid status of inpatients was tested positive prior to enrolment or during hospitalisation. Then, based on that test, we discriminated 306 patient as *Covid*+ and 78 patients as *Covid*− (see Fig.1a). In addition, we wanted to improve our models’ interpretability by means of the detailed clinical outcomes available from each patient at day 28 of their stay in the hospital. In total, those variables encompass up to 40 different types of multi-variated sequences (see *variable descriptions* file in SI). Those data allowed us to compute a precise overall-score of severity based on discrete World Health Organisation (WHO) Organisation cited November 2021 scores provided in the cohort for each patient over time of stay in the hospital. First, we vectorised those scores with the aim of being using entropy Gray 2013 to calculate the information encapsulated by the WHO scores in each patient. That information in bits Murphy 2012 enabled the construction of a probability density function that basically well stratified individuals according to severity progression of disease what ultimately reinforced the explanatory power of our learning models.

**Figure 1.**
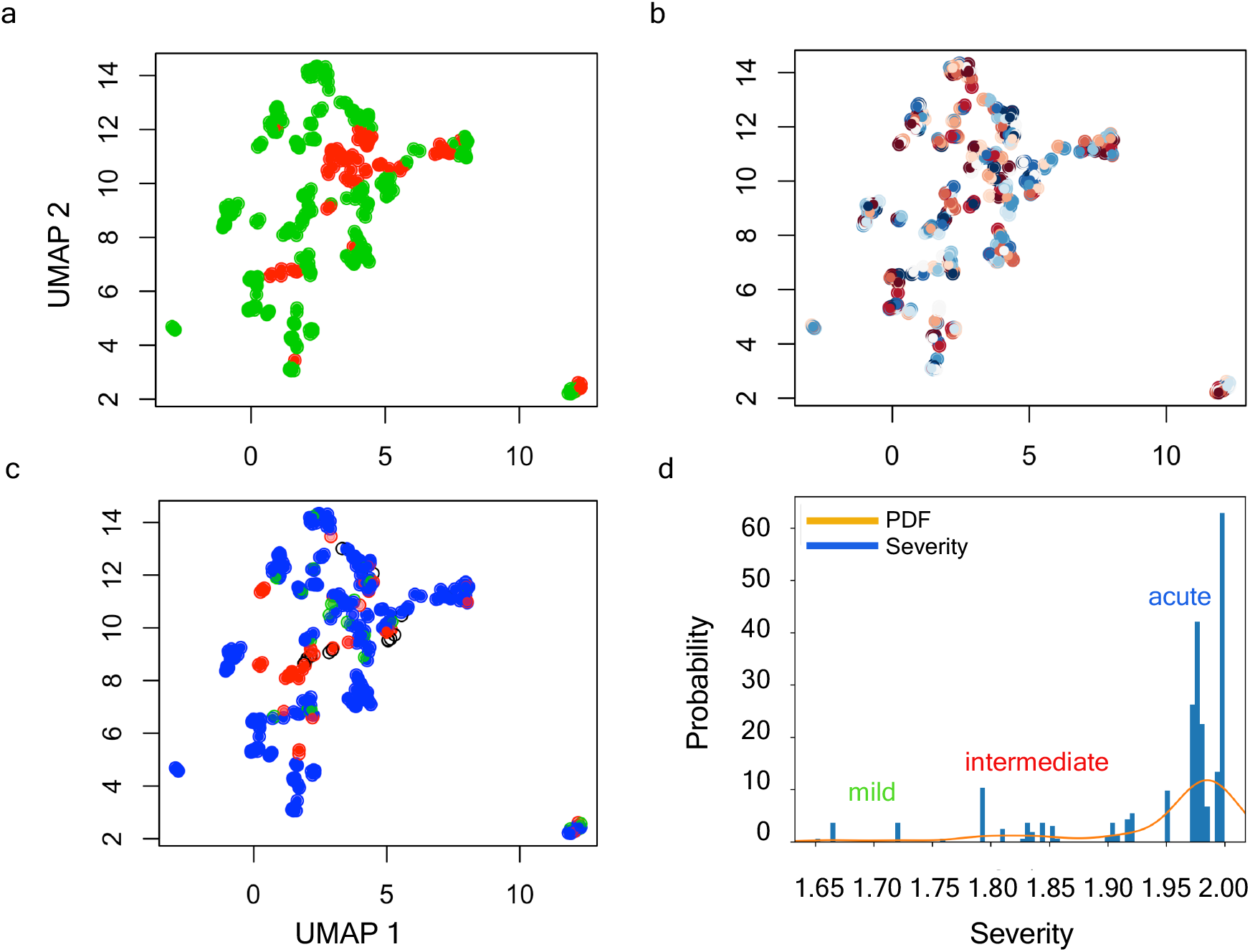
Hdbscan Severity clustering by means of Entropy measures. (a-c) UMAP projection of clinical data. (a) Stratification by *Covid*+ and *Covid*. (b) Inpatients’ stratification by WHO_*max*_ score (i.e., a measurement correlated with the maximum WHO outcomes achieved by patients during their hospital stay and also available in the clinical dataset). (c) Individual discrimination by Shannon’s Entropy combined with *Hdbscan* clustering algorithm. The blank circles show inpatients considered as outliers by the dissimilarity of their symptoms. (d) Probability density function optimally fitted in accordance with Shannon’s Entropy displaying three sharp peaks, namely: mild, intermediate, and acute associated with inpatients of the *Covid* cohort.

## Results

### Hierarchical models of WHO scale-based entropy information stratify patients by severity

The scores of severity provided by WHO monitoring disease progression throughout the entire patient’s stay at the hospital centre. Those values were recorded according to the discrete measure *W*_*s*_ = {1, 2, …, 6} ∈ ℤ^+^ at days 0, 3, 7, and 28 (see *variable descriptions* file in SI for their interpretation). This is a suitable *clinical* indicator, but pretty local screenshot of disease proression in an individual (see Fig.1b). To avoid such an inconvenient, we intended to construct a way of containing local and global records along with some meaningful qualitative information. We considered, then, information theory scores by Shannon 2001. Thus, we vectorised patient’s WHO scores obtaining a discrete random variable *V*, with possible outcomes *v*_1_, …, *v*_6_, which occur with probability *P* (*v*_1_), …, *P* (*v*_6_). In order to extract the self-information of any given event 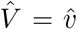 in bits MacKay 2017, we computed the entropy of each individual per vector by applying the formula 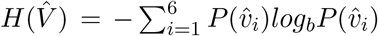 (see Fig.1c). Then, the bits information for a context channel 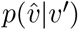 with 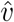 and *v*^*′*^ in the discrete alphabet *W*_*s*_ resulted in a continuous distribution whose domain 𝒟 = [1, 2] ∈ ℝ^+^. Hence, we fitted the transformed data (see Fig.S1a) to the best distribution by checking a comprehensive set of probability density functions (PDFs) (see Fig.S1b). The generalised continuous normal random variable (see Fig.1d) yielded the best performance and was calculated as follows:

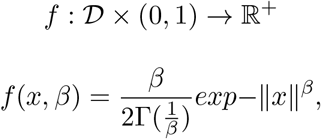

where *β* ∈ [0.01, 0.99] and Γ is the function gamma Sun 2020. By means of this function, we calibrated hierarchical clustering models that were computed applying the Hdbscan algorithm Campello, Moulavi, and Sander 2013. Hence, we achieved to discriminate the Covid cohort into three different groups (see files *mild/intermediate/acute_severity_Covidp* files in SI). Those groups basically met the *mild, intermediate*, and *acute* symptoms as registered in the available clinical dataset (see Fig.1d). From this stratification, 13 out of 384 patients were excluded to be considered as outliers with noisy data. These patients displayed dissimilar symptoms and unmatched characteristics amongst them to be included in any of the groups (see file *ids0_noisy_severity_cluster_Covidp* in SI and Fig.S2).

### The latent space of clinical features explains patients’ stratification

We unified many local perspectives of the clinical dataset to explain models of severity progression (see Fig.S4a-d). To this end, we summarised both an entire model and individual features learnt from the pdf of our Shannon’s Entropy severity. This task was eventually performed using the medical outcome dataset (see *MGH_COVID_Clinical_Info* file in SI) to train a three dense layers convolutional neural network with 269, 313 trainable parameters (see Fig.S3). The architecture of this network consisted of two convolution 2*D* and a last flatten layer with a scheme of (384, 40, 512) × 2 and (384 ∗ (40//512) ∗ 1) as output dimension. Next, we computed local explanations based on Shapley-related extensions, i.e., the so-called SHAP values Lundberg and Lee 2017. To figure out the relative contribution of each feature to our model output individually, we plotted the values of every local feature for every sample in the cohort. The Fig.2d -right hand panel-show a plot of sorted features by the sum of local value magnitudes over all samples, and uses such values to show the distribution of the impacts each feature has on the model output.

The colour represents the feature value (red high, blue low). This reveals for example, a known fact, that a high *categorical age* (% lower status of the population) raises the predicted risk of experiencing acute severity in Covid disease. In the left-hand panel of Fig.2d, we observe this same effect in stacked red bars for our multi-class output task.

**Figure 2.**
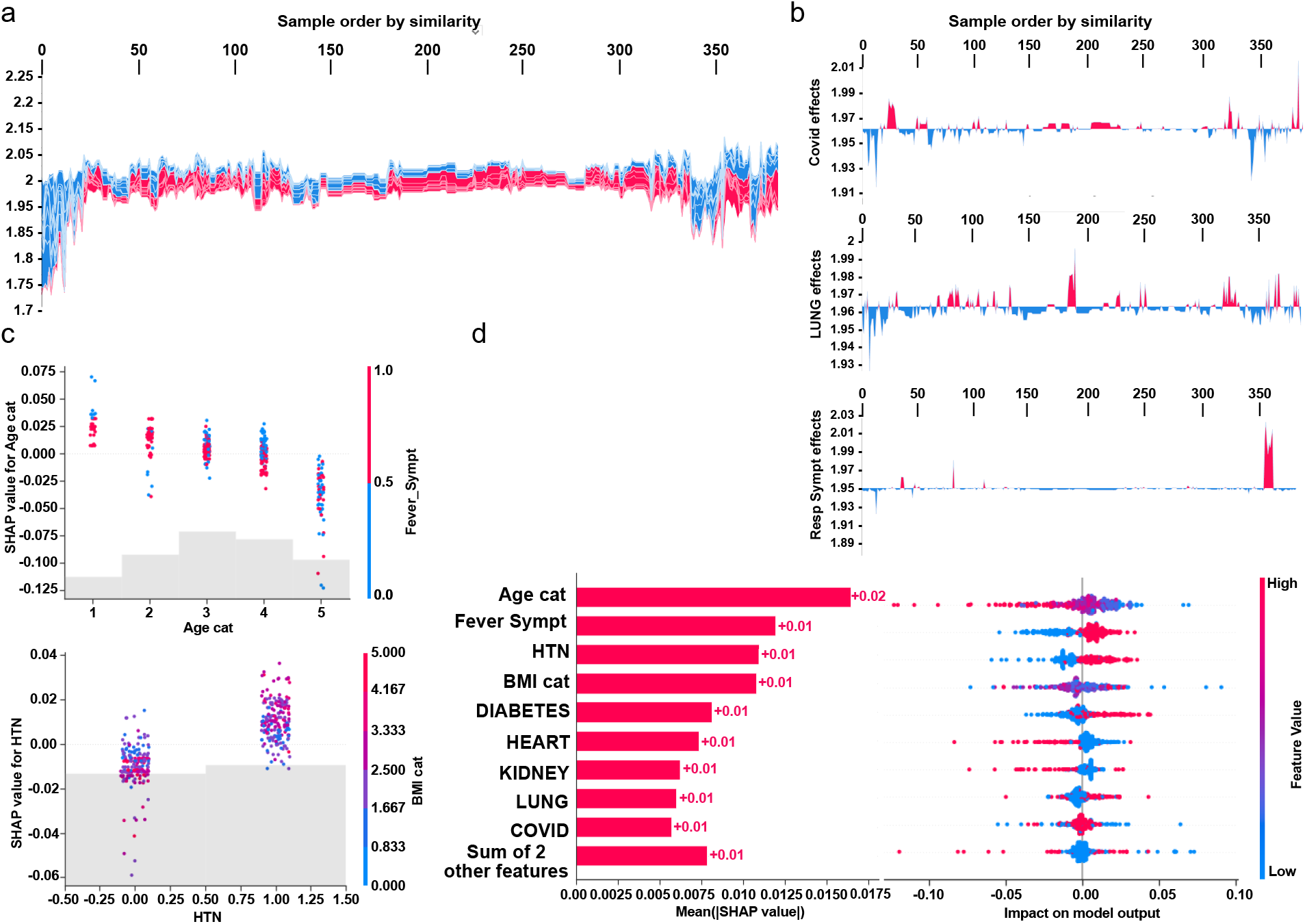
Clinical evaluation of our entropy-based model on Covid severity score by model and higher individual features. Initial explanations are based on a gradient boosted decision tree model trained on the covid cohort. (a) Globally stacked *SHAP* explanations clustered by explanation similarity. Inpatient profiles land on the *x*-axis. Red values increase the model prediction, blue ones decrease it. Two clusters stands out: On the left is a group with low predicted risk of suffering an acute covid, whereas on the right we have a group with a high predicted risk of suffering from acute covid. (b) Top-bottom: locally stacked explanations clustered by explanation similarity for infection, lung, and respiratory symptoms. (c) Effect of a single feature across the whole cohort. Top-bottom: dependence plots for *Age* and *Hypertension*(*HTN*) features. These plots display inflection points in predicted age and hypertension as *Agecat* and *HTN* (oldness by years on average and hypertension complaint per individual in the cohort) changes. Vertical dispersion at a single category of *Age* (resp. *HTN*) represents interaction effects with other features. To help reveal these interactions, we coloured by *Fever* (resp. *BMI*). We passed the whole explanation tensor to the colour argument in the dependence plots to pick the best feature to colour by. In this case, it selected Fever symptoms (resp. Body Mass Index) since that highlights that the average age (hypertension) per inpatient has more (less) impact on covid severity for categories with a low (high) *Fever* (*BMI*) value. (d) Left: bar chart of the average *SHAP* value magnitude. *Age* was the most important symptom, changing the predicted absolute covid probability on average by 2 percentage points (0.02 on *x*-axis). Right: a set of beeswarm plots, where each dot corresponds to an inpatient in the cohort per significative symptom. The dot’s position on the *x* axis shows the impact that a symptom has on the model’s prediction for a given inpatient. The piled up dots mean density of inpatients suffering from a symptom with similar impact on the model. Younger ages reduce the predicted covid risk, elder ages increase the risk.

To understand how a single feature effects the output of the model we plotted the local value of that feature vs. the value of the feature for all the inpatients in the clinical dataset (see Fig.2a). Now, we can zoom in some of those effects individually as shown in Fig.2b for lung, and respiratory (the one quantified as lowest in its contribution to severity learning) symptoms. Since locally explained values represent a feature’s responsibility for a change in the model output, the plot in Fig.2c represents the change in predicted Covid severity as *Age cat* (the average age per category in the cohort), or pre-existing hypertension change. Vertical dispersion at a single value of *Age cat* represents interaction effects with other features. To help reveal these interactions we can colour by another feature. If we pass the whole explanation tensor to the colour argument the scatter plot will pick the best feature to colour by. In this case it picks *Fever_Sympt* (symptoms associated with fever) since that highlights that the the average age per category in the cohort has less impact on acute Covid severity for categories with a high *Fever_Sympt* value.

The values of interaction between locally explained variables are a generalisation of those to higher order interactions. Fast exact computation of pairwise interactions are implemented for tree models with. This returns a matrix for every prediction, where the main effects are on the diagonal and the interaction effects are off-diagonal. These values often reveal interesting hidden relationships, such as how the increased risk of death peaks for inpatients with mild febrile symptoms at the age between 20 and 34 (see Fig.2c -upper panel-). Or that non-pre-existing hypertension has less impact on individuals with a high *BMI _cat* value (see Fig.2c -lower panel-).

### Persistent homology identifies novel key proteomic features involved in severity

Based on the previous clinical characterisation of Covid severity, we exploited the proteomic plasma information available for the remaining 371 individuals in the cohort. Unfortunately, the particular geometry of inpatient’s proteomes as embedded onto lower dimensional spaces resulted highly sensitive to parameter setups considered in downstream analyses according to entropy-based severity. In such scenario, we computed topological invariant structures instead (see *rips_ complex_patient_29_severe* video in SI). These invariants, the so-called simplicial complex, qualitatively analyse features that persist across multiple scales. Such invariants can be classified over days 0, 3, and 7 by obtaining their generators through persistent homology (see Fig. S5a). This analysis led us to identify unique protein configurations (see Figs.S5b and S6) within inpatient proteomes based on their connected components Aktas, Akbas, and Fatmaoui 2019; Xia and Wei 2014. Thus, the whole universe of proteome embeddings could be enclosed in the quotient space 𝒫/ ∼ under the given equivalent relation: for any *p*_•_ ∈ 𝒫, *p*_*i*_ ∼ *p*_*j*_ if the projected proteome *i* “is similar to” on the set of all rotated tails, the so-called special orthogonal group *SO*(3) Hall 2004. Hence, a partition with two classes of equivalence come out whose disjoint union determines the all three revealed groups of patients (see *rips complex triang patients 29 and 42 severe* video in SI). Indeed, these classes of equivalence have as represents [*c*_*b*_] := {*x* ∈ 𝒫 : *x* ∼ *c*_*b*_} iff *b* is a ball-like shape in *SO*(3) and [*c*_*s*_] := {*x* ∈ 𝒫 : *x* ∼ *c*_*s*_} iff *s* is a start-like shape in *SO*(3). These two classes can be visualise in upper and lower panels of Fig.3a-b. Now, to identify proteins whose profiles are invariants of each inpatient over the days of their stay {0, 3, 7}, we primary analysed the classes of equivalence [*c*_*b*_] and [*c*_*s*_] of each partition by persistent homology per group. Specifically, we used persistence diagrams wherein we quantified the number of homology generators (see upper Fig.3c) while testing their quality by means of confidence band generated by probabilistic boosting and density difussion (see lower panel of Fig.3c). This enabled the identification of unique set of proteins (see files *-M1s_mild / M2s_intermediate/ M3s_acute-_severity_Covidp* in SI) that encapsulated two and three dimensional structures (Figs.3d-e) important to topologically characterise severity classification per group over the days of stay of each individual in the hospital. Thus, we mapped the transmembrane serine proteases TMPRSS5 and SS15 Huttlin et al. 2017 as novel receptors taking part of the infection machinery. These proteins belong to the same family of TMPRSS2, a known receptor used by Sars-CoV-2 to enter the cell Hoffmann et al. 2020. Both of those proteins were located amongst the dysregulated interactome of inpatients stratified as acute and strikingly also as mild. Furthermore, amid the proteins we found (see *M#s* files in SI for the entire list), there were proteins in acute -BRK1, LAP3, SLC27A4, SLC39A14-, intermediate -SLC27A4-, and mild -BRK1, SLC27A4, and SLC39A14-inpatients functionally linked Huan, Sherman, and Lempicki 2009a,b with AP3B1, BRD4, BRD2, CWC27, SLC44A2, and ZC3H18 whose profiles are reported to be likely involved in early infection caused by the virus Gordon et al. 2020. Finally, the functional analysis of the novel proteomic features resulted from our persistence analysis showed along with their ancestors two sharp clusters bound to pneumonia and inflammation pathways (see Fig.S7).

**Figure 3.**
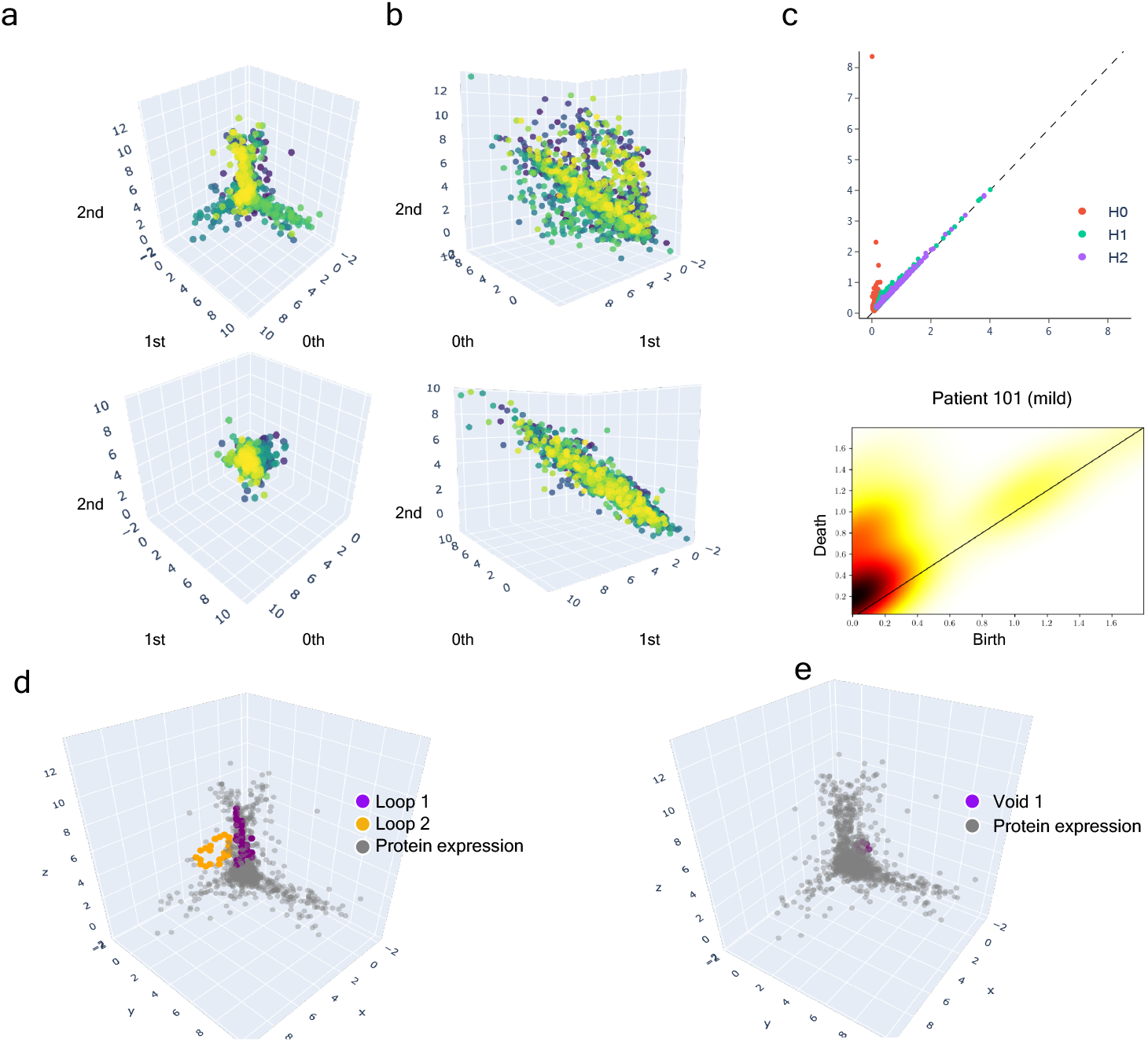
Multi-scale topology analysis flowchart. (a) -Upper-class of equivalence [*c*_*b*_] determined by umap projection of mild inpatients upon rotation on *SO*(3). -Lower- umap projection of an inpatient’s soluble proteome. (b) -Upper-class of equivalence [*c*_*t*_], taking as example to show the mild inpatient 101. -Lower-umap projection of that inpatient’s soluble proteome. (c) -Upper-topological feature extraction from diagram of persistence of patient 101 and its later calibration. -Lower-Application of density diffusion for separating noise from robust signals in persistence diagram of impatient 101. (d) Spotted loops of proteins enclosing dimension 2 structures important to explain severity stratification over time of patient 101. (e) Spotted voids of proteins enclosing dimension 3 structures important to explain severity stratification over time of patient 101.

### Dynamic tracking of protein interactions required by the virus to efficiently infect the cell

Once we put the spotlight on individual proteins topologically important to discriminate Covid patients over time, we envisaged to capture their dynamics of functional interactions at regulatory levels. To this end, integrated protein-protein interaction networks were firstly constructed per group of patients using co-localization, co-expression, physical interactions, and shared domains Warde-Farley et al. 2010. Then, we enquired these graphs about their connections quality by means of degree and centrality distributions as shown in Fig.4. Surprisingly, we could confirm (see Fig.4) that neither was highly connected nor played an important modular role in the graphs. Next, we monitored the behavioural regulation of the nodes’ graph aggregation on semi-supervised learning on a community composed by known and unknown protein interaction with ACE2 and TMPRSS2 Morilla et al. 2022 (see the movies in format .avi provided in SI, namely: *mild / intermediate / acute_fcD (resp. outlier)_severity_covidp_candidates_anim*.). To compute such a tracking, we endowed the graphs with a tailored hybrid design covered in convolutional layers along with a spectral rule Defferrard, Bresson, and Vandergheynst 2017 as occurs in graph convolutional networks (GCNs). The first model’s performance yielded an accuracy, for each group of inpatients, of 0.49, 0.41, and 0.85 supported by 76, 146, and 355 samples regarding *covid* and *non* – *covid* feature representations, respectively. Those values raised to 0.71, 0.84, and 0.95 to the second learning model. We also found their corresponding performances asymptotically tended to 0.65, 0.7, and 0.81 when layers were largely increased from the 32 units in the convolutional architecture.

**Figure 4.**
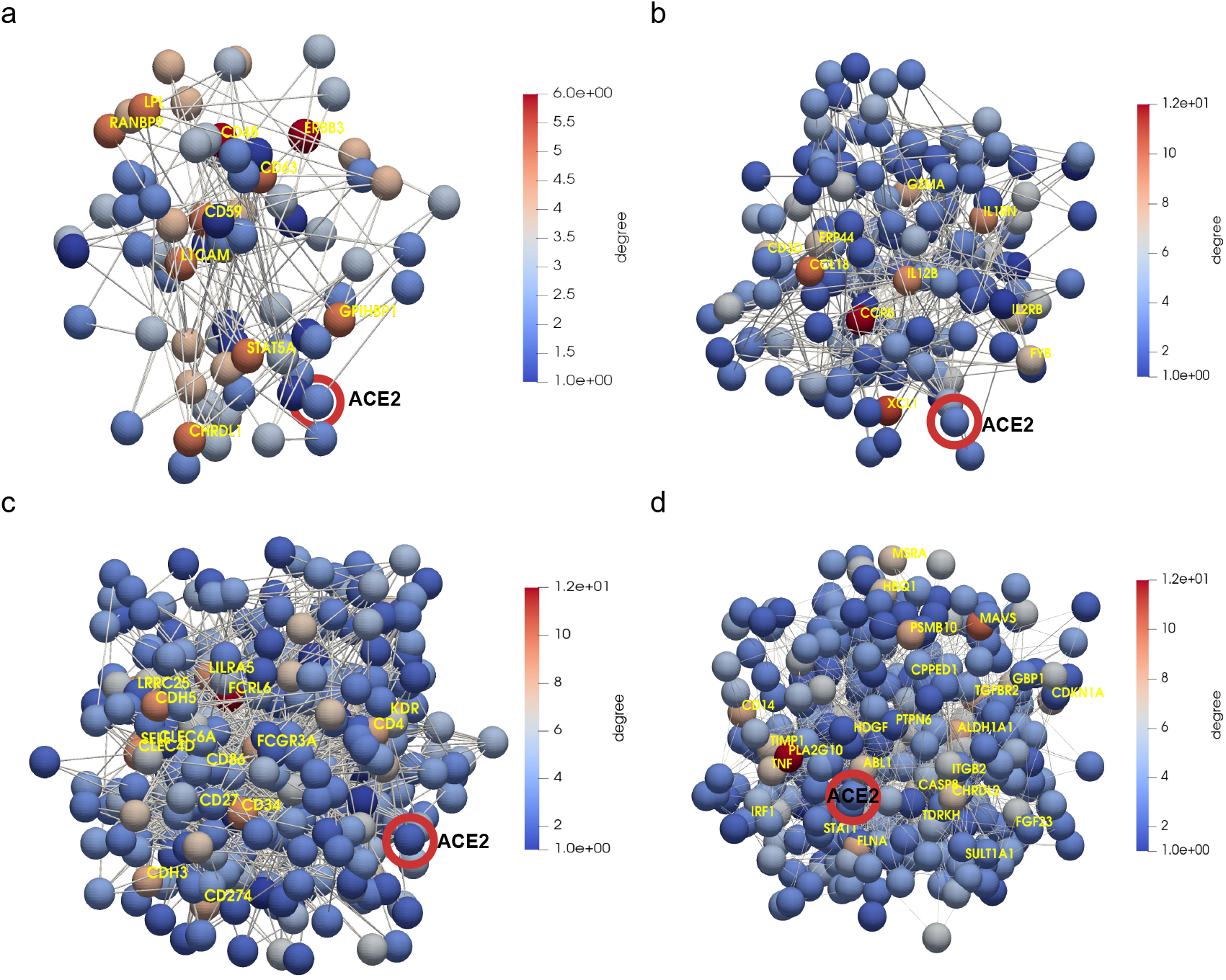
Regulatory gene networks regarding ACE2 and TMPRSS2. (a) mild patients. (b) intermediate patients. (c) acute patients. (d) group of proteins non-functionally enriched in acute patients. Highlighted in red ACE2 as one initial seem in the downstream analysis of protein interactions occurred post-infection. Yellow enhances those proteins (ERBB3, CD48, CCR5, FCRL6, PLA2G10, amongst others) with a higher connectivity degree in the networks.

In that way, we learnt how ACE2 and TMPRSS2 interacted with the persistent novel candidates to explain the virus machinery at its entrance into the cell to put patients into mild, intermediate or acute groups of severity over time (see mp4 video files *covidp_mild / intermediate / acute_feature_ # _weighted* in SI). Hence, a primary set of proteins that leaded to acute severity consisted of the progressive aggregation of BCAN, CA2, CA12, CLEC4, FOLR1, FOLR2, IFNGR2, IGSF3 (R), ILR13A1(R), LAIR1, LRRN1, PCDH17, RTBDN, SEZ6L, SIGLEC6 Schulte-Schrepping et al. 2020, and TNFRSF21 with respect to *covid* feature representation. Especially, CLEC4 belongs to a protein family (i.e., the C-type lectin receptor) involved in regulating immune reactivity through platelet degranulation whose expression showed significantly decreased in COVID-19 and correlated with disease severity Overmyer et al. 2021. The interactions occurring early on during the infection amongst AXL, CD58, DDR1, DLK1, FCGR3A, TNFRSF12A, UXS1, and XPNPEP2 set the *non*−*covid* latent feature. Herein, we spotted CD58 a nonclassical monocyte such as CD274 (PD-L1) known inhibitor of T cell activation along with Arginase 1 (ARG1) Bronte et al. 2003; Li et al. 2018 highly expressed in neutrophils in COVID-19 patients or CD24 involved in neutrophil degranulation with an increased expression of neutrophil function Overmyer et al. 2021. Therein, we also identified FCGR3A (encoding CD16a) that is regulating severity-dependent alterations of the myeloid cell compartment during Sars-CoV-2 infection. Indeed, FCGR3A has been already found to be a non-classical monocytes marker in COVID-19 Schulte-Schrepping et al. 2020. Next, to the *covid* representations that determined intermediate severity of patients, we found the early aggregations of FR2, GALS4, IL1RN(R), ILR1(R), LRPAP1, RNF41, TRIM21(R), and VWV2. Remarkably, RNF41 plays a central role during interactions of Sars-CoV-2 with innate immune pathways since its interferon pathway is targeted by RNF41 (NSP15) Gordon et al. 2020. Then, the intermediate *non* − *covid* representations are governed by the interactions of CCR5 (intestinal pro-inflammatory), CPA1, LAMA2, PLA2G4A, PON3, SETMAR, TGFB1, and XCL1. Finally, aggregations of C4BPB, CD70(R), IPCEF1, MAVS(R), PLCG2, and THBS2(R), leaded to mild severity stratification to *covid* feature representations. At the same time, the interactions between CD200, MAPKAPK5 Kindrachuk and al 2015, NTRK3(R), PRAP1(R), XPNPEP2(R) set the *non* − *covid* feature representations. In these two list, we might mention a similar effect on neutrophils and expression as CD58 to CD70 and CD200 Overmyer et al. 2021.

We checked that *covid* feature representation of patients with acute symptoms was functionally charaterised by a set of proteins involved in the fusion of virus enclosure to the host endosome membrane (GO:0039654) at the virus entrance into the cell. Overall, the interactions between ACE2 and TMPRSS2 and these persistent proteins were largely enriched in the immunoglobulin-like fold functional category. From a mere clinical stratification point of view, these conditions characterise most of the inpatients in the available clinical dataset suffering of cardiovascular complications. *Non* − *covid* representation of acute patients was strongly composed by membrane and signal peptide functional categories, protein tyrosine kinase and glycoprotein with extracellular and cytoplasmatic topological domains and transmembrane helix and integral component in the region biological processes. In this case, these conditions felt on patients mainly suffering from diabetes of type 1 and immunosuppression.

Regarding the *covid* feature representation of intermediate patients, an overabundance of protein binding function is observed with diabetes of type 2 and normal variation diseases associated with such condition. On the other side, the *non* − *covid* feature of this group is strongly enriched with disulfide bond and signal peptide functional categories. Therein, we found various terms directly linked with endosome viruses’ machinery. Thus, we identified clathrin-dependent endocytosis (GO:0075512), host lysis, inhibition of host IKBKE, JAK1, RLR pathway, TBK1, and TLR pathway triggered by virus in the host cell. Those symptoms were identified to a wide range of the complications described in the clinical dataset. In particular, asthma severity, chronic hepatitis C, immunosuppression after liver transplantation, diabetes specially strong of type 2, heart and kidney complications, and hypertension.

Finally, the latent *covid* representation of mild patients’ proteomes were functionally characterised by a weak overabundance of disulfide bond and glycosylation site (i.e., N-linked as GlcNAc, etc.). These categories were related with supression by virus of host adaptative immune response (GO:0039504). Remarkably, there were no disease-associated genes type-specific to these biological processes. The *non* − *covid* mild features were actually overrepresented by signal peptide, qualitatively similar to those features described to the intermediate *non* − *covid* patients. We will fully expose and discuss the intriguing implications of such results in the next section.

## Discussion

Sars-CoV-2 has become these two last years a real life-thread that has collapsed the health systems worldwide. Many efforts have been already done to structurally characterise the Sars-CoV-2 spike protein. To predict its severity, large mappings of proteins likely involved in the machinery applied by the virus to infect the cell have been reported. All these investigations have led to enormous advancements in COVID-19 treatment that consequently have given rise to efficient vaccines. However, there is still some facets not well-characterised or yet sufficiently explored. In our attempt to contribute to this research, we computed an overall severity score based on WHO scales instrumental to provide a chart explaining the protein interactions required by the virus to stratify a patient’s infection into mild, intermediate, or acute. To this end, we made use of a double analysis linking symptoms to protein expressions and interactions with ACE2 and TMPRSS2. Merely from a stratification standpoint, we give novel information as well as verified known facts about Covid. As conclusion, we could claim that in itself COVID-19 is not as harmful as it is in association with other risk factors such as age, febrile symptoms, or overweight. Indeed, most of the *Covid*− patients, though some were considered as outliers as indicated in the earlier sections, but most held a relative high entropy value of severity due to an eventual intubation, ventilation or supplementary oxygen requirement. To achieve those high peaks *Covid*+ should be elderly people with overweight and present fever and/or respiratory symptoms.

More importantly, we obtained functional evidences of how particular sequences of proteins interacted with the virus to block the immune systems during the infection. Thus, we could explain the infection fate towards the acute symptoms since an endosome aciditication is produced during the infection initiating conformational proteins fusion. In that way, Sars-CoV-2 could take advantage of such pathway to be endocytosed as happens with many types of viruses such as influenza A virus, alphaviruses or HIV-1.

Regarding patients suffering from intermediate symptoms that endocytic process is caused over proteins that contain at least one coiled domain forming stiff bundles of fibres. Hence, proteins are modified by signalling upon creation of interchain disulfide bonds, which can produce stable, covalently linked protein complexes likely contributing to fold and stabilise proteins. In virus internalization, clathrin-mediated endocytosis could be then generated in response getting assembled on the inside face of the cell membrane to cleave the host cell (CCV) by the action of DNM1/Dynamin-1 or DNM2/Dynamin-2. Then, the virus may be delivering their content to early endosomes via CCV. These mechanisms could be expressing in Sars-CoV-2 using different ways as by lysing the host cell, blocking the host innate defenses via IKBKE/IKK-epsilon kinase inhibition, JAK1 protein, DDX58/RIG-I-like repector (RLR) what stabilises the antiviral state, TBK1 kinase inhibition to prevent IRFs activation, or toll-like recognition receptor (TLR) pathway evasion, which makes the production of interferons to be inhibited and so to establish a stable antiviral state.

To mild cases, the Sars-CoV-2 protein could be preventing the tuned repertoire of self and nonself antigens’ recognition of efficiently acting against the malicious effects of cell infection. In these cases, Sars-CoV-2 would be escaping the adaptive immune response by simply interference with the presentation of antigenic peptides at the surface of infected cells.

Overall, results provided in this work, contribute to gaining new insights into non-linear relationship between “message passing” proteins, particularly explaining disease severity modulation during the Sars-CoV-2 infection.

## Methods

### Samples

Data provided by the MGH Emergency Department COVID-19 with Olink proteomic. Clinical dataset and plasma proteomes of 384 patients distributed in 306 patient that tested positive in COVID-19 that were named as *Covid*+ and 78 patients tagged as *Covid*− that tested negative even suffering from respiratory symptoms.

### Notes on the graph convolutional network

We endowed the regulatory persistent-based protein networks (RPPNs) with a convolutional design (i.e. neural networks) along with a spectral rule of node aggregation as in GCN Defferrard, Bresson, and Vandergheynst 2017. The sequential combination of two hybrid models enabled the learning of interactions between ACE2 and TMPRSS2, and the persistent proteins needed by the virus to be spread in cells. Following this reasoning, we primary made use of the identity matrix *I* as features and the adjacency matrix *A* contributing the model in the following spectral rule:

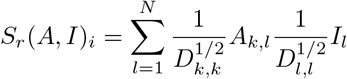

where *D* is the degree matrix. Right after, we considered the metric given by distance of the shortest path to characterise the early aggregation of persistent proteins to ACE2 and TMPRSS2 as an additional feature in a second model. Thus, we generated models with two layers computed by 32 units per layer and a 2D transformation of the activation function tanh. When applying the spectral rule the relu activation function is applied at the beginning of the layer implementation instead of later on. The number of epochs was set to 250 and 5000, respectively. We computed the stochastic gradient descent (sgd) optimizer in the training task picking a learning rate and momentum regularization set to 0.001 and flagged true, respectively. The semisupervised classification (*covid* or *non* − *covid* early proteins aggregation) of nodes in the RPPNs Kipf and Welling 2017 was performed by an in-house python script based on MXNet implementation *Faster*, Cheaper, Leaner: improving real-time ML inference using Apache MXNet 2021.

## Data access

All data used in this work are included in the article and/or SI Appendix.

Acknowledgments

We would like to thanks the funding from National Research Association (ANR) (Inflamex renewal 10-LABX-0017 to I.M.), DHU FIRE Emergence 4, and the l’Agence de la Biomédecine.

